# Phenotypic diversity created by a transposable element increases productivity and resistance to competitors in plant populations

**DOI:** 10.1101/2021.10.04.462998

**Authors:** Vít Latzel, Javier Puy, Michael Thieme, Etienne Bucher, Lars Götzenberger, Francesco de Bello

## Abstract

An accumulating body of evidence indicates that natural plant populations harbour a large diversity of transposable elements (TEs). TEs provide genetic and epigenetic variation that can substantially translate into changes in plant phenotypes. Despite the wealth of data on the ecological and evolutionary effects of TEs on plant individuals, we have virtually no information on the role of TEs on populations and ecosystem functioning. On the example of *Arabidopsis thaliana*, we demonstrate that TE-generated variation creates differentiation in ecologically important functional traits. In particular, we show that *Arabidopsis* populations with increasing diversity of individuals differing in copy numbers of the *ONSEN* retrotransposon had higher phenotypic and functional diversity. Moreover, increased diversity enhanced population productivity and reduced performance of interspecific competitors. We conclude that TE-generated diversity can have similar effects on ecosystem as usually documented for other biological diversity effects.

## Introduction

It has been progressively manifested that biological diversity is one of the key determinants of ecosystem functioning (Huston 1997; Loreau & Hector 2001). Diversity effects have been mostly assessed at the level of species diversity, mostly number of species (Tilman et al. 1997; Díaz et al. 2007), and interpreted as the effect of phenotypic differences between species, i.e. greater functional trait diversity, enhancing ecosystem functioning (Díaz et al. 2007; Gross et al. 2017). In species diversity studies, positive diversity effects imply that communities composed by phenotypically different organisms are expected to be more productive, stable and resistant to abiotic and biotic stresses such as competition from potential invaders, compared to less diverse ones (Gross et al. 2007). This positive diversity effects on ecosystem functioning could be because increasing species richness increases the likelihood that one or a few species with a large effect on any given ecosystem property would be present (i.e. the so-called section or sampling effect; Loreau & Hector 2001); and because greater niche differences between coexisting species result in a better utilization of resources by the community (i.e. so-called complementarity; Loreau & Hector 2001).

Important evidence has been accumulating that also phenotypic diversity within populations can have strong and positive effects on ecosystem functioning comparable to the one of species diversity (Crutsinger et al. 2006; Ehlers et al. 2016). This phenotypic variation within populations has been most often attributed to genetic variation (Zhu et al. 2000; Hughes et al. 2008; Kotowska et al. 2010; Zuppinger-Dingley et al. 2014), originating from ongoing spontaneous genetic mutations and recombination, migrations from surrounding populations (Ulukapi & Nasircilar 2018) as well as due to epigenetic processes (Latzel et al. 2013; Puy et al. 2020). Additionally, at the crossroad between genetic and epigenetic diversity, variation in transposable elements (i.e. mobile genetic elements that can replicate and/or change position within genomes; Slotkin & Martienssen2007) represent a mechanism that can change DNA sequence variation as well as epigenetic regulation of gene expression and thus generate phenotypic variation (Kidwell and Lish 1997; Niu et al. 2018). However, it remains unclear whether variation in transposable elements within populations cause sufficient phenotypic variation to substantially affect ecosystem functionality.

Transposable elements (TEs, also called “jumping genes”) are a ubiquitous component of DNA of most eukaryotes (Feschotte et al. 2002), making up the majority of DNA in case of some plant species (Bennetzen et al. 2005; Schnable et al. 2009). TEs are commonly divided in two major classes: DNA-transposons and retrotransposons. While DNA-TEs (Class II-TEs) move by a “cut-and-paste” mechanism, retrotransposons (Class I-TEs) replicate (i.e. multiply) via an RNA intermediate with a “copy-and-paste” strategy (Wicker et. al. 2007). The mutational power of unrestricted TE-mobility can result in genomic instability and lead to negative effects on host function. To protect their genome, plants have evolved complex silencing mechanisms such as RNA-directed DNA-methylation to restrict their activity (Matzke and Mosher 2014). Because TEs attract epigenetic regulation such as DNA-methylation and histone modifications (Sigman and Slotkin 2016) to their insertions site and also act as transcriptional regulators (Butelli 2012), transposition events can substantially alter host gene expression in plants (Lopez & Bureau 2014). Thus, TE-induced phenotypic diversity can not only be explained by genetics but also be based on altered epigenetic regulation (McClintock 1984; Dubin et al. 2018; Domínguez et al. 2020).

Importantly, the phenotypic variation generated by TEs can arise over very short time scales, even within one or few generations due to heritable transpositional bursts (Feschotte et al. 2002). Transpositional bursts are large numbers of simultaneous transpositions of TEs, which are triggered by challenging situations including genomic or environmental stress (McClintock 1984; Wessler 1996; Kidwell & Lish 1997; Maumus et al. 2009; Ito et al. 2011; Rey et al. 2016). Hence, even originally genetically (quasi-) uniform populations (e.g. newly established small populations from limited diaspora, selfing or clonal individuals) can potentially develop substantial and rapid phenotypic variability due to stress-induced mobilisation of TEs (e.g. Belyayev et al. 2010; Domínguez et al. 2020). Overall, TEs can contribute to increase the phenotypic variation, in other words, enhance the diversity of functional traits of the population (i.e. functional diversity).

While the effect of TEs on plant phenotypic variation has been studied almost exclusively at the level of individuals (either from ecological or evolutionary perspective), there is scarce information on the effect of TEs on the functioning of plant populations or whole ecosystems. Theoretically, such intraspecific functional diversity in a population could have a similar effect to that produced by interspecific functional diversity, i.e. species diversity in a community (Loreau & Hector 2001; Cadotte 2017). In a within-population context, mixtures of individuals differing in the number and positions of TEs in DNA, by increasing the functional diversity of the population, could be more productive and resistant than monocultures consisting of individuals with identical number and positions of TEs. Functional diversity mechanisms, though common in plant populations, are unexplored from the TEs perspective. It is thus highly desirable to test whether TE-generated diversity affects population’s functioning similarly to species diversity or genetic diversity effects.

In our study, we took advantage of available *Arabidopsis thaliana* lines (further referred to as TE lines) created by Thieme et al. (2017a). TE lines were derived from the Col-0 accession by stress induced mobilization of the endogenous *ONSEN* (AtCOPIA78) retrotransposon (heat-responsive copia-like retrotransposon, Cavrak et al. 2014; Ito et al. 2011; more details on TE line creation are provided in Methods or in Thieme et al. 2017a,b). Importantly, the novel *ONSEN*-copies are stable in their positions in DNA, heritable over at least three generations and create phenotypic variation (Thieme et al. 2017a,b). From the original pool, we used 18 unique TE lines plus 2 control lines for which we characterized a number of functional traits important for species coexistence and ecosystem functioning. We constructed populations that differed in the number of TE lines: populations were composed from one TE line (‘monocultures’), or from two, four, or sixteen different TE lines (further referred to as ‘mixtures’). We subjected designed mixtures and monoculture populations to different abiotic and biotic conditions (control, drought, interspecific competition or combination of drought and competition). This design allowed us answering whether there is a substantial effect of TEs diversity on productivity and resistance of plant populations to abiotic and biotic stress. Specifically, we tested (T1) to which extent TE lines differed between them in terms of key functional traits associated with plant fitness and effects on ecosystem functions; (T2) whether an increasing number of TE lines increased functional diversity of populations and enhanced their functioning (increased *Arabidopsis* productivity and decreased performance of interspecific competitors) and whether this effect was consistent across abiotic conditions (changes in water regimes). The effects of functional diversity can be associated to niche complementarity effects (Díaz et al. 2007; Cadotte 2017). Finally, we also tested (T3) whether diversity, either estimated by number of TE lines or functional diversity, affected population functioning beside the effect of specific TE lines and their characteristics (the latter corresponding to the so-called ‘selection effect’; Loreau & Hector 2001).

## Material and Methods

### Plant material: TE lines

TE lines were derived from *A. thaliana* Col-0 accession that was germinated and transiently grown on a medium containing zebularine and α-amanitin. Zebularine is a Cytidine analogue known to inhibit DNA-methyltransferases in plants which leads to a decrease of DNA-methylation resulting in de-repression of TEs (Baubec et al. 2009), while α-amanitin inhibits RNA polymerase II (Pol II), which has been suggested to repress the *ONSEN-TE* (Thieme et al. 2017a). Combination of the two chemicals together with exposure of seedlings to heat stress led to mobilisation of *ONSEN* that resulted in an increase of genomic *ONSEN* copies in DNA (Thieme et al. 2017a). Identifying several TE lines differing in the number of *ONSEN* copies was achieved by screening the F1 generation of heat stressed and drug-treated plants by qPCR and transposon display to identify and select high-copy lines (Thieme et al. 2017a,b). From the list of available TE lines we selected 20 lines with the number of *ONSEN* copies ranging from 8 to 60 (Thieme et al. 2017a, b). Two out of the 20 TE lines were controls, i.e. they did not undergo *ONSEN* mobilisation. Since the control lines did not deviate from the rest (see later, Fig. S1) and because *ONSEN* copies are present in their DNA too, for simplicity we further consider them as TE lines (TE lines 19 and 20).

### Biodiversity-population functioning experiment

The study was carried out in a heated greenhouse of the Institute of Botany in Průhonice from January to April. From the pool of 20 TE lines, we established populations of *A. thaliana* with differing diversity, achieved by manipulating the number of different lines sown in each population. Populations consisted of 48 seeds sown into pots (12 cm in diameter), either of the same TE line (i.e. monocultures) or mixtures of two TE lines (24 seeds per TE line), four TE lines (12 seeds per TE line) and sixteen TE lines (3 seeds per TE line). For the mixtures, for each diversity level, we selected TE lines randomly (random generation of combinations) with the condition that all 20 TE lines are equally represented (i.e. across populations within each treatment combination, see next). Further, all the populations were randomly assigned to one of the four following treatments that differ in the abiotic and biotic conditions: control – no manipulation, drought – population watered only if showing significant drought stress, i.e. leaves were wilting, competition – we planted 3 seeds of *Plantago lanceolata* and 15 seeds of *Poa annua* into the pot, drought + competition – we combined drought and competition described above.

The monoculture for each TE line was replicated four times per treatment (4 x 20 x 4=320), and the mixtures were replicated 20 times per treatment and diversity level (20 × 4 × 3 = 240). Thus, the final set-up comprised a total of 560 experimental populations from 26 880 sown seeds of *A. thaliana* that resulted in 15 390 harvested individuals twelve weeks after establishment of the experiment. At the end of the experiment, we harvested above ground biomass in all pots and recorded the number of *A. thaliana* individuals in each population. The biomass of *A. thaliana* and, when present, both competitors (*Plantago lanceolata* and *Poa annua*) were separated, dried at 60° C for 48 h and weighed. *Arabidopsis’* biomass and competitors’ biomass (of each pot) were used as estimates of population functioning: (a) *Arabidopsis* total productivity and (b) performance of the competitors respectively.

The lowest germination rate was around 65% for some lines (determined in a germination trial prior the population’s construction) suggesting that at least one individual per each line germinated in each mixture. The number of harvested individuals of *A. thaliana* in monocultures showed that the lowest establishment rate was 35% of sown individuals of TE-line 1 and the highest establishment was 78% of sown individuals of TE-line 12, F=5.77, p<0.0001. We used standardised soil substrate mixed with sand in a 1:1 volume ratio. We distributed seeds randomly and evenly on the soil surface using two layers of tiny mesh. All pots were placed in a cold chamber room (4° C) for 4 days for stratification before moving them to the heated greenhouse. In the greenhouse, we set the day regime to 9-10 hours (ambient light conditions) during the first month and to a long day regime of 14 h for the rest of the experiment (artificial light supplemented in the morning and late afternoon). The mean temperature was maintained at 23/18° day/night.

### Functional traits

To approximate the phenotypic variation of the 20 selected TE lines, we set up a parallel study to the main experiment where we screened phenotypes of each of the lines and measured a number of plant functional traits known to affect organism’s fitness, species coexistence and ecosystem functions (de Bello et al. 2010; Adler et al. 2014). This separate experiment was set-up because the measurements were destructive. In this experiment we grew 10 replicate plants of each line (10 replicates x 20 lines, 200 plants in total) alone in the controlled environment of a growth chamber (Fitotron® Plant Growth Chamber) set to an 8 hours day with temperature regime 22°/18° C day/night. We used the same substrate as in the main study. We changed positions of all pots on a weekly basis. After 5 weeks from germination, we measured the total dry biomass of each individual (including both aboveground plus root biomass), and several above- and belowground vegetative traits connected with the so-called ‘plant economic spectrum’ (Reich 2014; Díaz et al. 2016), reflecting resource use strategies in plants. For each plant, one leave was scanned to estimate the leaf area and weighed for fresh mass and dry mass after drying at 60° C (48h). We used these measurements to estimate specific leaf area (SLA; leaf area per dry mass, mm^2^/mg) and leaf dry matter content (LDMC; leaf dry mass per leaf fresh mass, mg/mg). In addition, roots were carefully extracted, washed and scanned at 1200 dpi with an Epson Perfection 4990 scanner. From the scans, total root length, average root diameter (mm), and the distribution of root length per different diameter classes were determined by using the image analysis software WinRHIZO Pro, 2008 (Regent Instruments Inc., Quebec, Canada). After scanning, roots were dried for 48 h at 60 °C and weighed. We used these measurements to estimate root biomass allocation (i.e. root mass factor; RMF; root biomass per total biomass, g/g), specific root length (SRL; root length per dry mass, m/g) and percentage of very fine roots (roots with a diameter < 0.2mm from the total).

### Data analysis

#### Phenotypic differences across TE-lines (T1)

To assess phenotypic differences across TE-lines (T1) we followed two approaches. First, using the 10 replicate plants for each TE line, we tested the independent variation of each single trait (total biomass, RMF, SLA, LDMC, SRL, and % of fine roots) across lines, with line identity as predictor in a one-way ANOVA. Tukey post-hoc tests were used to verify significant differences between lines (Fig. S1). Second, we tested trait differences across all traits together. This was done by computing a multivariate trait dissimilarity between each pair of the 200 individual plants, using Euclidean distance based on scaled trait values (i.e. values were centred by the mean and divided by 1 standard deviation unit for each trait). Then we used the multivariate trait dissimilarity as a response variable and line identity as predictor in a PERMANOVA (Anderson 2001), which is the corresponding multivariate approach to univariate ANOVA although in this case the significance value is tested by permutations.

#### Diversity effect on population functioning (T2)

### Number of TE-lines diversity effect

We applied a number of different approaches to assess the effect of the diversity of TE lines on productivity and competitors’ performance. First, we assessed the effect of the number of TE lines in the population (i.e. 1, 2, 4, 16) on productivity (total *Arabidopsis’* population dry biomass) and on the performance of the competitors (dry biomass of *Plantago lanceolata* and *Poa annua*). To compare productivity across treatments and TE diversity (i.e. number of TE-lines) we fitted a linear model with treatment (4 levels: control, drought, competition and drought + competition) and (log-transformed) number of lines (1, 2, 4, and 16), and their interaction as predictors of productivity. A significant interaction term will indicate that diversity effect on population functioning (i.e. productivity and resistance to competitors) are different across treatments. Since the effect of treatments was strongly prevailing over the effect of diversity (either number of TE lines or functional diversity) and the interaction term between treatment and diversity was not significant, we adopted a specific strategy to represent results, for graphical purposes: we used a model in which we focused on the effect of diversity after accounting for the effects of treatments (using the residuals of a model with treatments as only predictor, i.e. its partial effects). Differences between diversity levels were tested using pairwise t-tests. Similarly, identical models were also run for competitors’ performance as response variable, with only two treatments considered: i.e. competition with and without drought.

### Functional trait diversity effect

Then, we assessed the effect of the functional diversity resulting from mixing TE lines on productivity and competitors’ performance. To do so, similar models to the ones described in the previous section were employed: using treatments and functional diversity, instead of the number of TE lines, as predictors (i.e. the same test, but with functional diversity instead of number of TE lines in the model) to explain both productivity and competitors’ performance. The following steps were done to compute functional trait differences between TE-lines within each pot and compute functional diversity. We first computed for each measured trait its average for the 10 replicates for each of the 20 TE lines. Following this step, we used a Principal Component Analysis, PCA, to inspect the main axes of trait variation across TE lines, which also helped visualizing the main correlations between traits (the highest correlation was Pearson R=0.75 between SLA and total biomass; for all results, see Fig S2). Using trait averages per TE line we then computed a mean trait dissimilarity between each pair of TE lines, as explained above (Euclidean distance based on scaled trait values). Based on these values we could compute an average trait dissimilarity between TE lines for each pot, using the commonly used Rao index of functional trait diversity (de Bello et al. 2016). The index provides zero dissimilarity for monocultures (as the dissimilarity within a TE line is considered to be zero). The effect of functional diversity can be interpreted as a biodiversity effect attributed to niche complementarity (Diaz et al. 2007; Cadotte 2017).

#### Effect of specific TE lines and their traits on population functioning (T3)

Parallel to these analyses, we tested whether the presence of specific TE lines in the mixtures influenced *Arabidopsis* productivity and competitor’s performance. We did this by first running a model selection (based on most parsimonious stepwise selection process using AIC’s model fit) with either *Arabidopsis* productivity or competitor’s performance as dependent variables and treatments plus the presence of the 20 lines as predictors. This led to the identification of few lines whose presence affected more strongly productivity or competitor performance. Following this step, we combined the models described for T2 (with treatments and diversity of number of TEs lines as predictors) including those specific TE lines whose presence had an effect on the functioning of populations (stepwise selection process using AIC’s). The effect of specific lines on the functioning of populations can be interpreted in this case a biodiversity effect due to selection effects (Loreau & Hector 2001).

Similarly, to test whether the specific functional trait of the TE lines considered, rather than their specific identity, influenced productivity and competitor performance (a functional trait-based approach) we run identical model selection including the population functional trait means (i.e. average value of each of the six functional traits per population) instead of the 20 lines as predictors. The improved predictions of these new models, i.e. comparing T3 against T2 in the ones using TE lines (Models 1, 3 and 5 in Table S1) and in the ones using a functional trait-based approach (Models 2, 4 and 6 in Table S1) separately; was assessed by checking changes in R^2^ between models, as well as the Akaike information criterion (AIC). A drop of 2 AIC points was considered to be an improved model.

## Results

### Phenotypic differences across TE-lines (T1)

The experimentally induced bursts of *ONSEN* TE elements in *A. thaliana* Col-0 ecotype caused phenotypic differences associated to important functional traits (T1). TE lines differed in almost all measured functional traits except LDMC (Fig S1). The highest variation was recorded for total plant biomass (aboveground and root biomass). Overall, TE lines differed in their multi-trait phenotypic variability (TE line identity explained 35% of the trait dissimilarity between individual plants; PERMANOVA R^2^ = 0.35, p = 0.001). The main differences between TE lines were associated to the so-called plant economic spectrum (SLA and LDMC, first axis of the PCA axis, explaining ~50% of trait variation between lines; Fig. 1).

**Figure 1:**
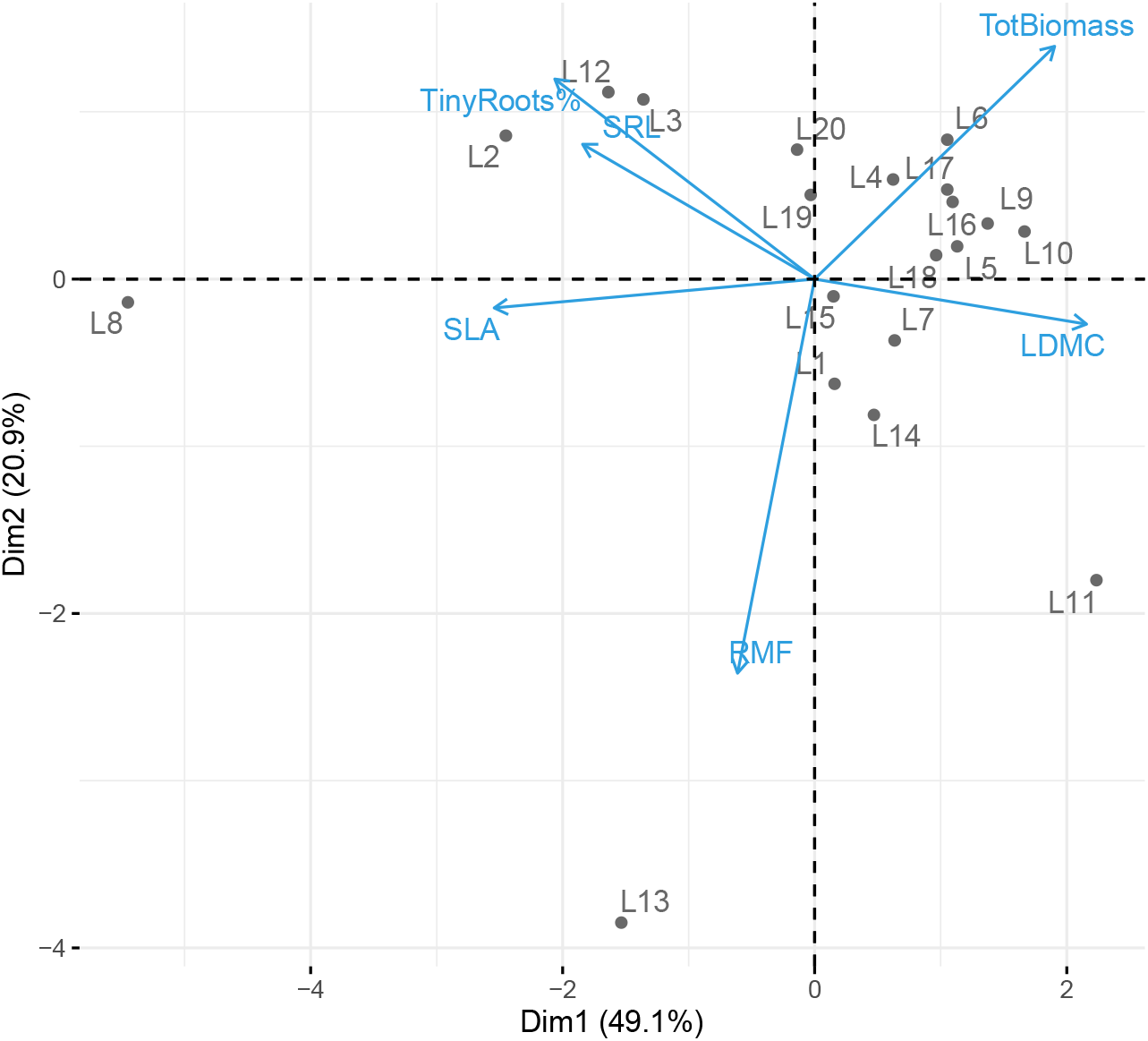
Principal component analysis (PCA) showing relationships among functional traits of TE lines. Blue arrows represent the traits used to build the principal component, i.e. total individual biomass (together aboveground and root biomass); specific leaf area (SLA), specific root length (SRL) and percentage of fine roots (% Tiny roots) which are typical traits indicative of a resource acquisitive strategy; and root mass factor (RMF), leaf dry matter content (LDMC) that are indicative of a resource conservative strategy.

### Diversity effect on population functioning (T2)

Compared to control conditions, all treatments (i.e., drought, competition and drought & competition) had a negative effect on productivity of *A. thaliana* populations (Fig. 2A; Model 1 in Table S1). The strongest reduction in productivity was observed in the treatment that combined biotic and abiotic stress: competition and drought. The performance of competitors (biomass of *Plantago lanceolata* and *Poa annua*) also decreased with drought (Fig. 2C) and decreased with increasing biomass of *A. thaliana* (Fig. S3). Mixtures of *Arabidopsis* TE lines were more productive than monocultures (Fig. 2B, pairwise t test: monocultures *vs*. mixtures p < 0.01), leading to positive overyielding for productivity (Fig. S4; i.e. lines in mixtures produced more biomass compared to when grown alone). This effect was irrespective of the treatment (no significant interaction treatment × number of TE lines; Model 1 in Table S1) although the effects tended to be slightly lower under drought conditions. Mixtures of *Arabidopsis* TE lines, compared to monocultures, also reduced the competitors’ performance (Fig. 2D; pairwise t test: monocultures *vs*. 2 TE lines mixtures p < 0.01, monocultures *vs*. 4 and 16 TE lines mixtures p = 0.06), and also irrespectively of drought treatment (no significant interaction treatment × number of TE lines; Model 1 in Table S1). Although mixtures of *Arabidopsis* TE lines were more productive and had lower performance of competitors than monocultures the increasing number of TE lines within the mixtures did not affect their functioning (i.e. neither productivity, either performance of competitors). In other words, mixtures of two TE-lines had comparable functioning to most diverse mixtures (Fig. 2B).

**Figure 2:**
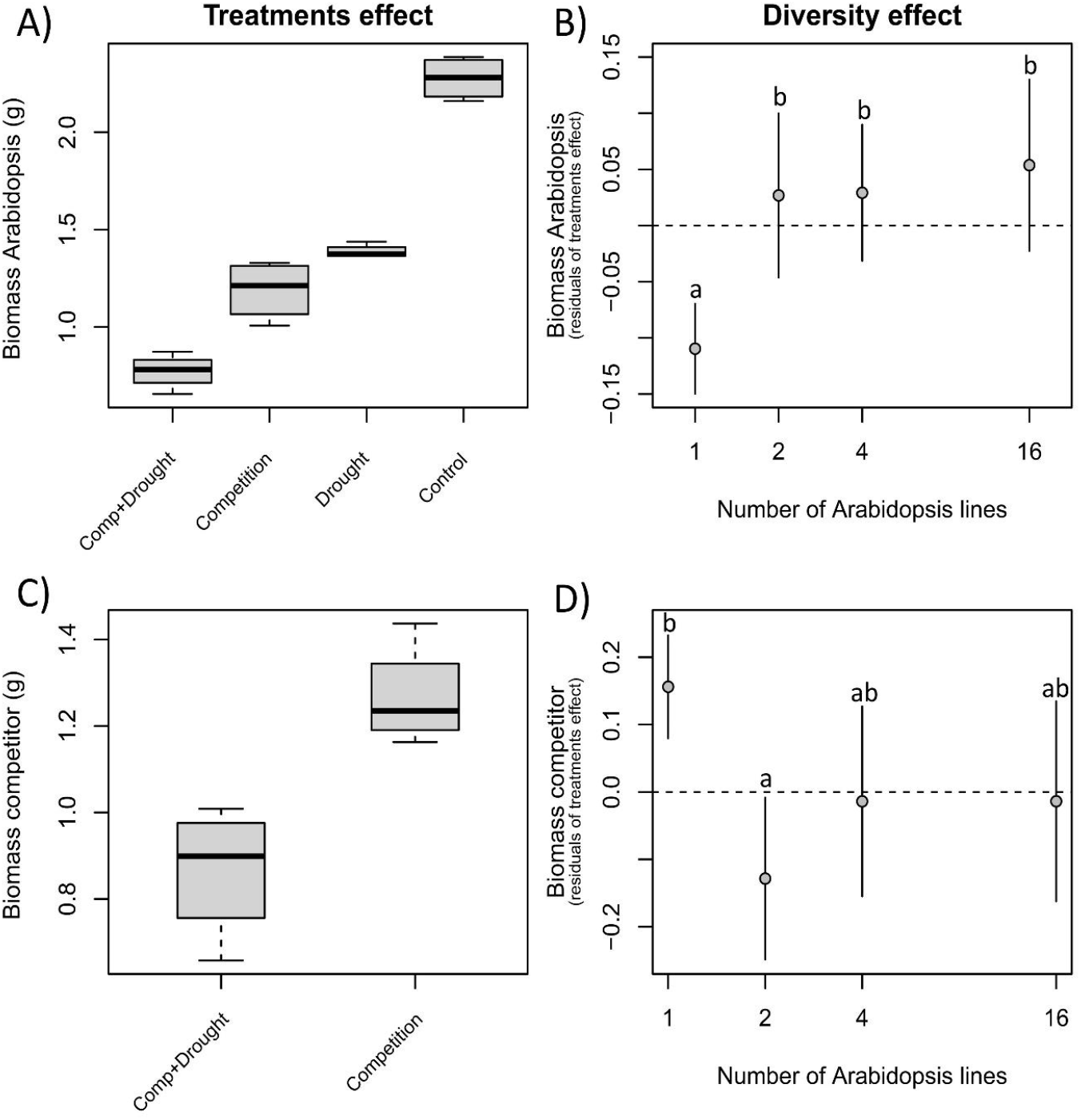
Effect of treatment (A and C) and diversity, measured as number of TE lines in the population (B and D; after accounting for the effects of treatments, its partial effects), on biomass of Arabidopsis (A and B) and biomass of the competitors (C and D). The bottom and top of the boxes are the 25th and 75th percentiles respectively, the centered band is the median and the whiskers represent 1.5 times the length of the box further from the box limits or the maximum or minimum observation in the absence of outliers. Dot and lines represent mean and 95% confidence intervals of each of the diversity levels. Different letters within each panel indicate significant differences between treatments (pairwise T test, p = .05)

As expected, phenotypic variability of populations increased with increasing numbers of TE lines (Pearson correlation between number of TE lines and functional diversity R^2^ = 0.67, p < 0.001 and increased mean functional diversity across the different number of TE lines, Fig. 3A). Due to this pattern, the effect of functional diversity on productivity and competitor performance (Fig. 3B, 3C) was comparable to the effect of the number of lines (Fig. 2B, 2D). In fact, using either the number of TE lines or functional diversity resulted in very similar model R^2^ (Models 1 *vs*. 2 in Table S1; for productivity: R^2^=0.77 using treatments and either number of lines or functional diversity; and for competitor’s performance R^2^=0.19 using number of TE lines as quantitative predictor and R^2^=0.21 using functional diversity).

**Figure 3:**
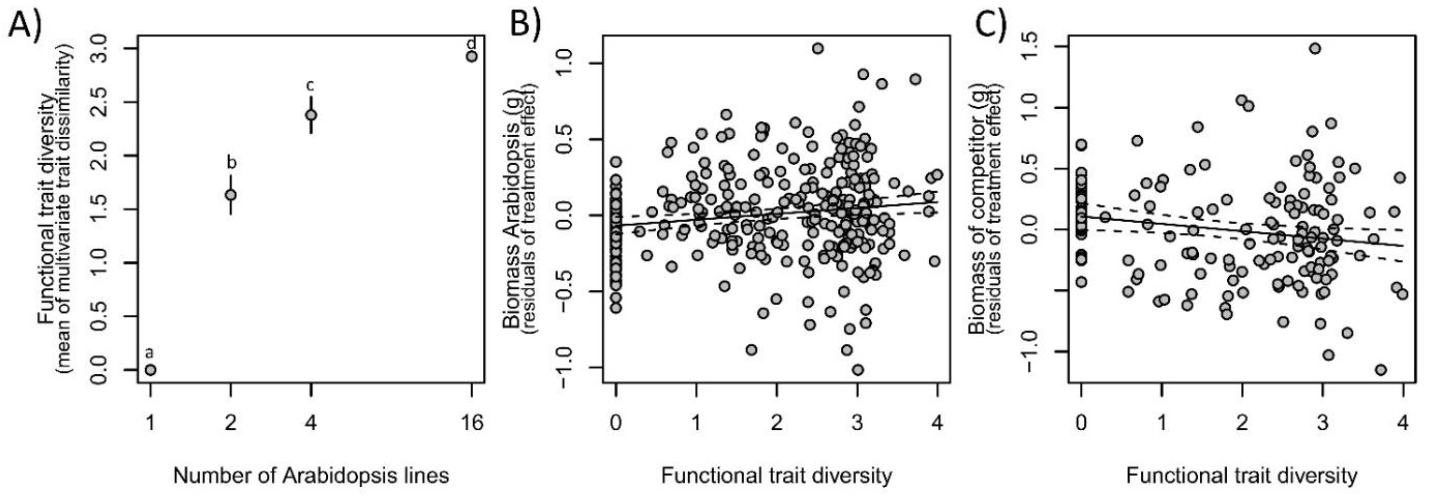
A) Corresponding functional diversity of each of the populations resulted from mixing differing number of TE lines. B) & C) Effect of diversity, measured as functional diversity (after accounting for the effects of treatments, its partial effects) on biomass of Arabidopsis (B) and biomass of the competitors (C). Dot and lines represent mean and 95% confidence intervals of each of the diversity levels. Different letters within each panel indicate significant differences between treatments (pairwise T test, p = .05). In regression plots, solid lines and dashed lines represent the slope estimate and 95% confidence intervals.

### Effect of specific TE lines and their traits on population functioning (T3)

We assessed whether individual TE lines or their specific traits, affect population performances by adding specific lines and population trait means to the models, alone and together with the corresponding diversity estimates (number of lines, or functional diversity, respectively). Generally, the inclusion of specific lines slightly improved predictions for both, *Arabidopsis* productivity and competitor’s performance (Models 3 and 5 in Table S1). Specifically, lines 8 and 15 had, respectively, a negative and positive significant effect on *Arabidopsis* productivity; and lines 2, 5 (positive) and 11 (negative) effect on the competitor’s performance (Model 5 in Table S1). The inclusion of functional traits of the populations, however, only improved predictions for *Arabidopsis* productivity (Models 4 and 6 in Table S1). *Arabidopsis* populations formed by TE lines with lower SLA tended to have more productivity, but competitors’ performance was not affected by the functional characteristics of *Arabidopsis* populations (Models 4 and 6 in Table S1).

## Discussion

We investigated the role of phenotypic variation of *Arabidopsis thaliana* populations created by experimental *ONSEN* retrotransposon bursts on generating intraspecific functional diversity and different effects on the functioning of populations. We show that the TE-generated variation can create variation in ecologically important functional traits such as plant biomass (both above and below-ground) or other traits associated with the so-called plant economic spectrum (e.g., SLA and SRL). Importantly, we showed that an increasing number of TE lines, i.e. lines with different number of *ONSEN* retrotransposon copies integrated at different places in the genome, increased functional diversity as well as the total population productivity. Moreover, higher functional diversity of *Arabidopsis* populations reduced the performance of their competitors, meaning that more diverse populations were more resistant to interspecific competitors.

Random transpositions of *ONSEN* retrotransposon in individual TE lines significantly influenced numerous functional traits associated with the plant economic spectrum (Reich 2014; Díaz et al. 2016) that reflect the resource-use strategies for plants. It is thus plausible that TE-generated variation translated into functional divergence among individuals for traits like SLA, SRL or biomass; and that this, by promoting niche and resource partitioning between TE lines, could have contributed to increased productivity of mixtures due to better utilization of the limited resources, i.e. due to a mechanism generally referred to as ‘complementarity’ (Hooper et al. 2005). Although the observed diversity effect seems to be mainly driven by functional diversity, pointing in the direction of complementarity (Díaz et al. 2007), we also identified specific TE lines and traits that had significant effects on productivity of mixtures. This indicates that the probability of including a key TE line in mixtures (specifically TE lines 8 and 15; Table S2), the so-called selection effect (Loreau & Hector 2001), also played a role in observed diversity effects. However, only characteristics of TE line 8 strongly deviated from the rest (i.e. smallest, and highest SLA; Fig S1), suggesting that although many traits linked to resource foraging strategy and competitive ability of the plants were measured, they were insufficient to completely characterize niche differences among TE lines (Kraft et al. 2015; Kunstler et al. 2016; Cadotte 2017). Nevertheless, we were able to detect an effect on population productivity and resistance to competitors by the average SLA of the population, which is a trait known to be related to resource foraging strategy of the plant (i.e. niche segregation), as well as to the fitness or competitive ability of individuals (Kraft *et al*., 2015; Puy et al. 2020). It is also interesting that the key TE lines in mixtures were the lines with higher *ONSEN* copies and not the original control lines (TE line 19 and 20) that did not undergo *ONSEN* mobilization. This means that *ONSEN* mobilization can generate new beneficial phenotypes that can be further selected over the original genotype.

Besides increasing population productivity, it is generally acknowledged that increased phenotypic diversity usually increases resistance of populations to environmental stress (e.g., Grime 2001 or Tilman et al. 2006). This was however not the case for our study as we did not observe a significant interaction between diversity level and treatment. This means that, although our treatments of contrasting abiotic and biotic environments (drought, competition, and both in combination) were clearly stressful for the populations of *Arabidopsis* (i.e. biomass production decreased by more than 50 % compared to control conditions), diversity did not ameliorate the negative effect of stressors when compared with monocultures. In other words, all populations were similarly affected by stressors despite their different levels of diversity. However, what we did find is that interspecific competitors produced less biomass in mixtures than when grown in *Arabidopsis* monocultures, irrespectively of whether mixtures were formed by 2, 4, or 16 TE lines. This decreased performance of competitors, although not translating into an ameliorating effect of competition on the productivity of *Arabidopsis* populations, may indicate that diversity enhances the resistance to invasion against competitors and population performance.

In nature, small and/or declining populations are often at risk of loss of genetic diversity due to inbreeding or random genetic drift. Low genetic diversity is inevitably connected with increased risk of populations’ extinction. Our study outlined that TEs can be beneficial to plant populations by creating functional diversity that promotes population functioning and stability in addition to generating heritable (epi)genetic variation on which selection (evolution) can act (Lisch 2009). This indicates that commonly reported maladaptive phenotypic consequences of TEs on individuals (e.g. Vinogradov 2003) can be, in particular situations, ameliorated or even overturned at the population level by the functional diversity effect. Moreover, considering that challenging situations including genomic or environmental stress trigger transpositional bursts, TEs can be seen as an insurance mechanism for populations to dramatically increase their functional diversity in response to stressors, allowing rapid adaptation to changing ecological conditions (Belyayev 2014). This mechanism might be especially crucial for the persistence of populations with limited dispersal capacity at the trailing edge of their distribution or suitability range. TE-generated functional diversity can thus provide the necessary time for adaptation in threatened populations, which ultimately can facilitate their survival.

To sum up, our proof-of-principle study indicates that TE generated functional diversity can affect functioning of populations similarly to effects that are otherwise documented for genetically diverse populations (Crutsinger et al. 2006; Ehlers et al. 2016).

## Acknowledgement

This study was financially supported by the Czech Science Foundation (GACR 20-13637S). JP was funded by the Irish Research Council Laureate Awards 2017/2018 (IRCLA/2017/60) to Yvonne Buckley. EB was funded by ERC Consolidator grant (BUNGEE 725701) of the European Union. FdB was supported by the Plan Nacional de I+D+i (project PGC2018-099027-B-I00), MT was financially supported by University Research Priority Programs (URPP) Evolution in Action and European Commission (PITN-GA-2013-608422–IDP BRIDGES).

## Author contributions

V.L. and J.P. designed and performed the study. F.B. and J.P. analysed the data. M.T. and E.B. provided plant material. All authors wrote the manuscript. V.L. and J.P. made equivalent contributions and should be considered joint first authors.

## Supplementary material

**Figure S1:**
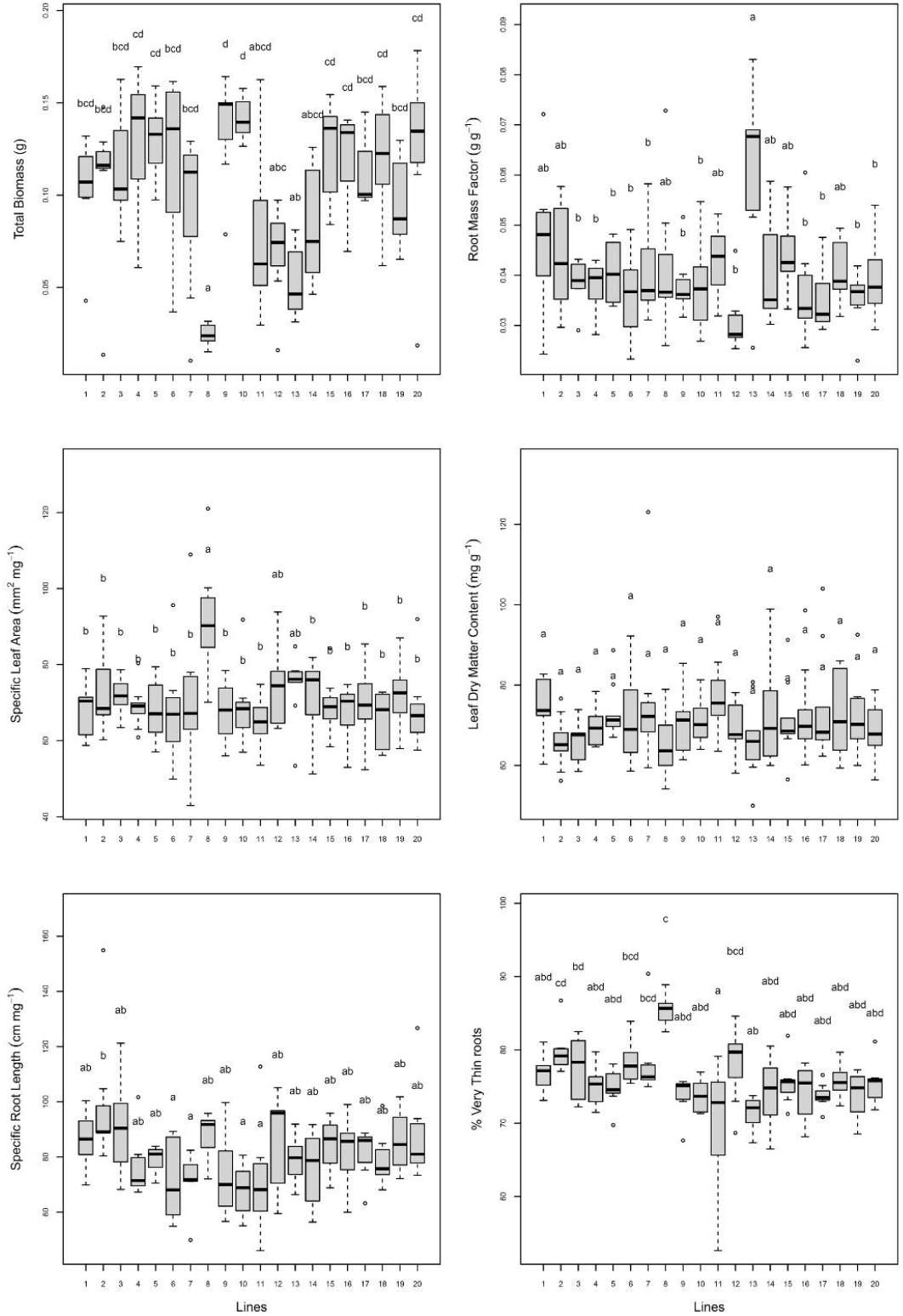
Difference between pairs of TE lines in the different functional traits measured. Each panel presents one of them: total individual biomass (together aboveground and root biomass); root mass factor (RMF); specific leaf area (SLA); leaf dry matter content (LDMC). specific root length (SRL) and percentage of fine roots (% Tiny roots). Different letters within each panel indicate significant differences between TE lines (Tuckey test, p = .05).

**Figure S2:**
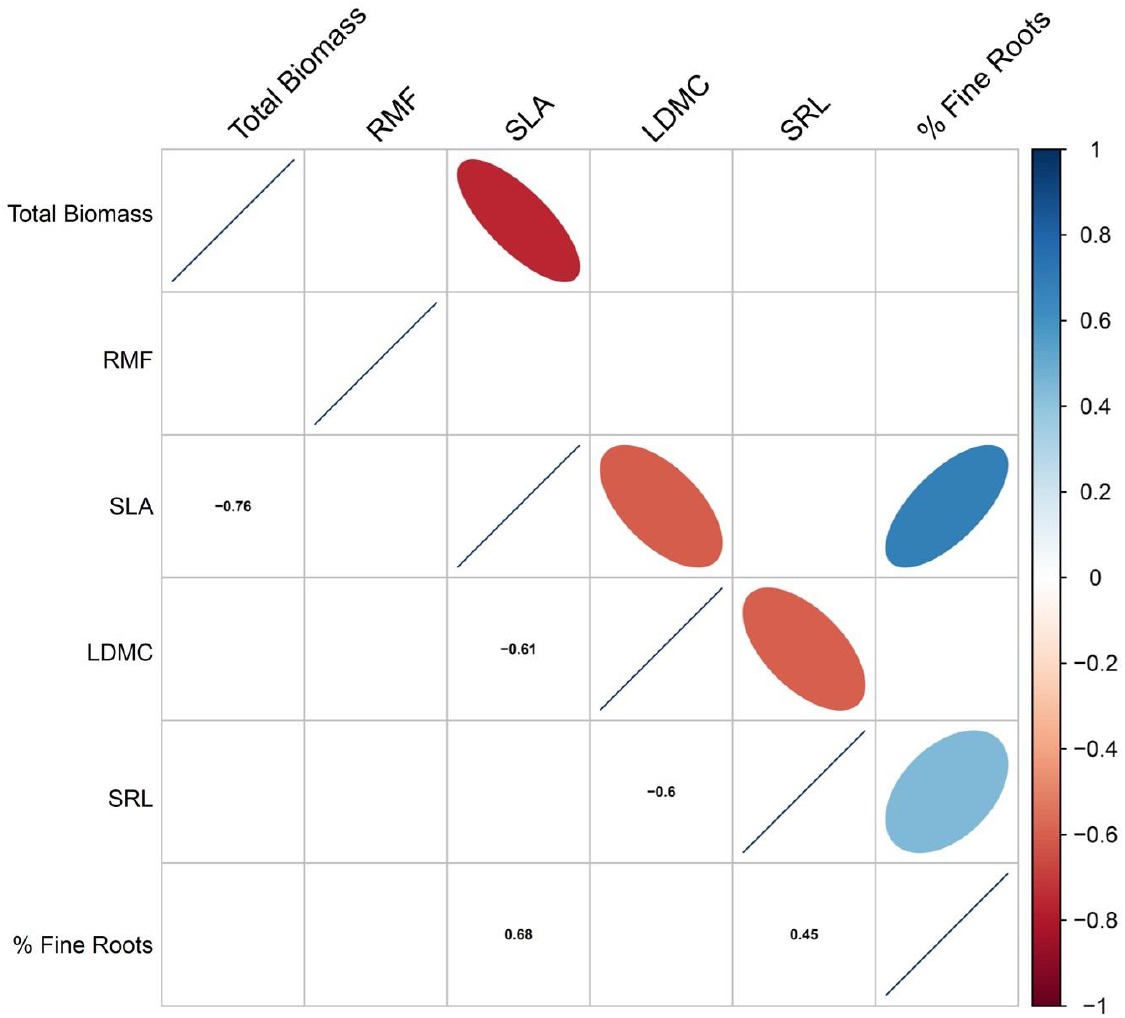
Correlation between pairs of functional traits measured in TE lines. Only significant correlations are represented. Below the diagonal, the numerical Pearson coefficients are displayed, while above the diagonal the coefficients are represented by coloured ellipses: blue is positive, red is negative, and the intensity of the colour represents the strength of the coefficient.

**Figure S3:**
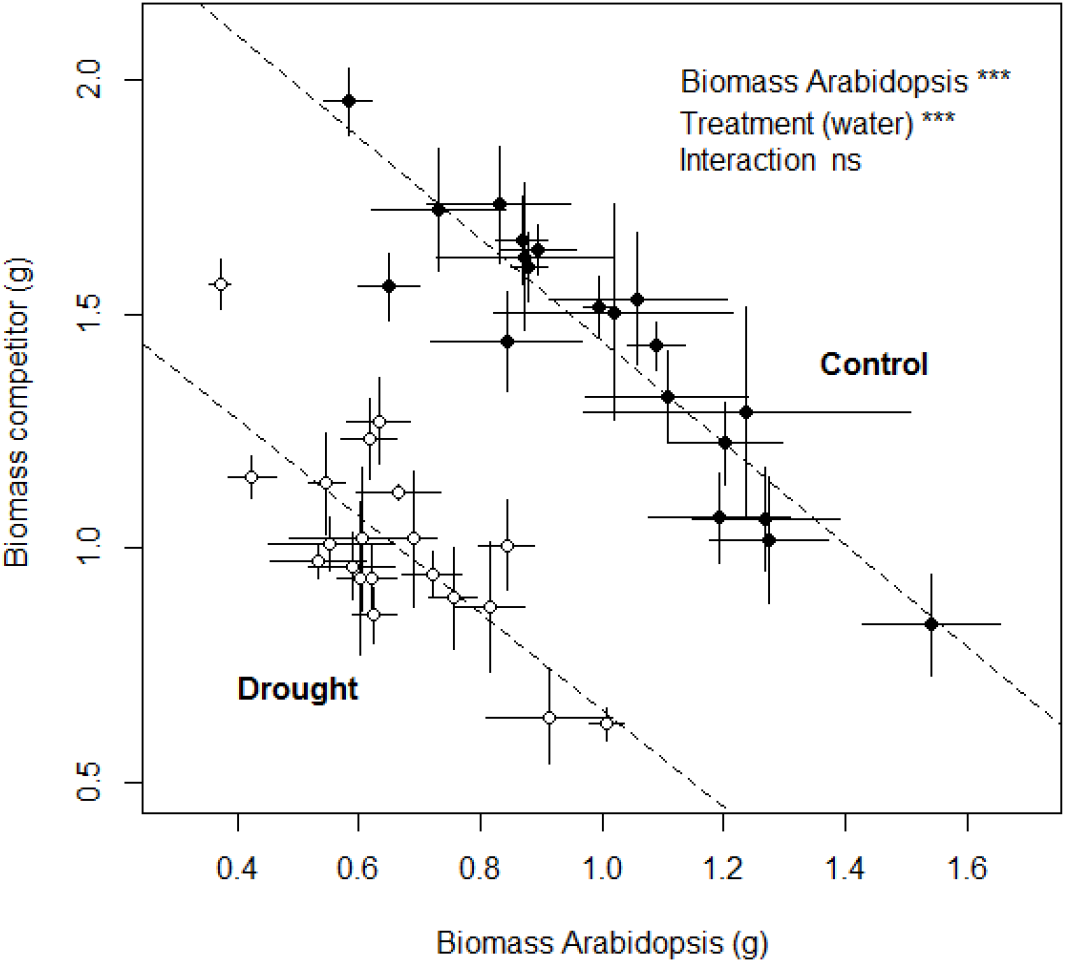
Effect of Arabidopsis biomass and Treatment (drought vs. control) on competitor’s performance.

**Fig S4:**
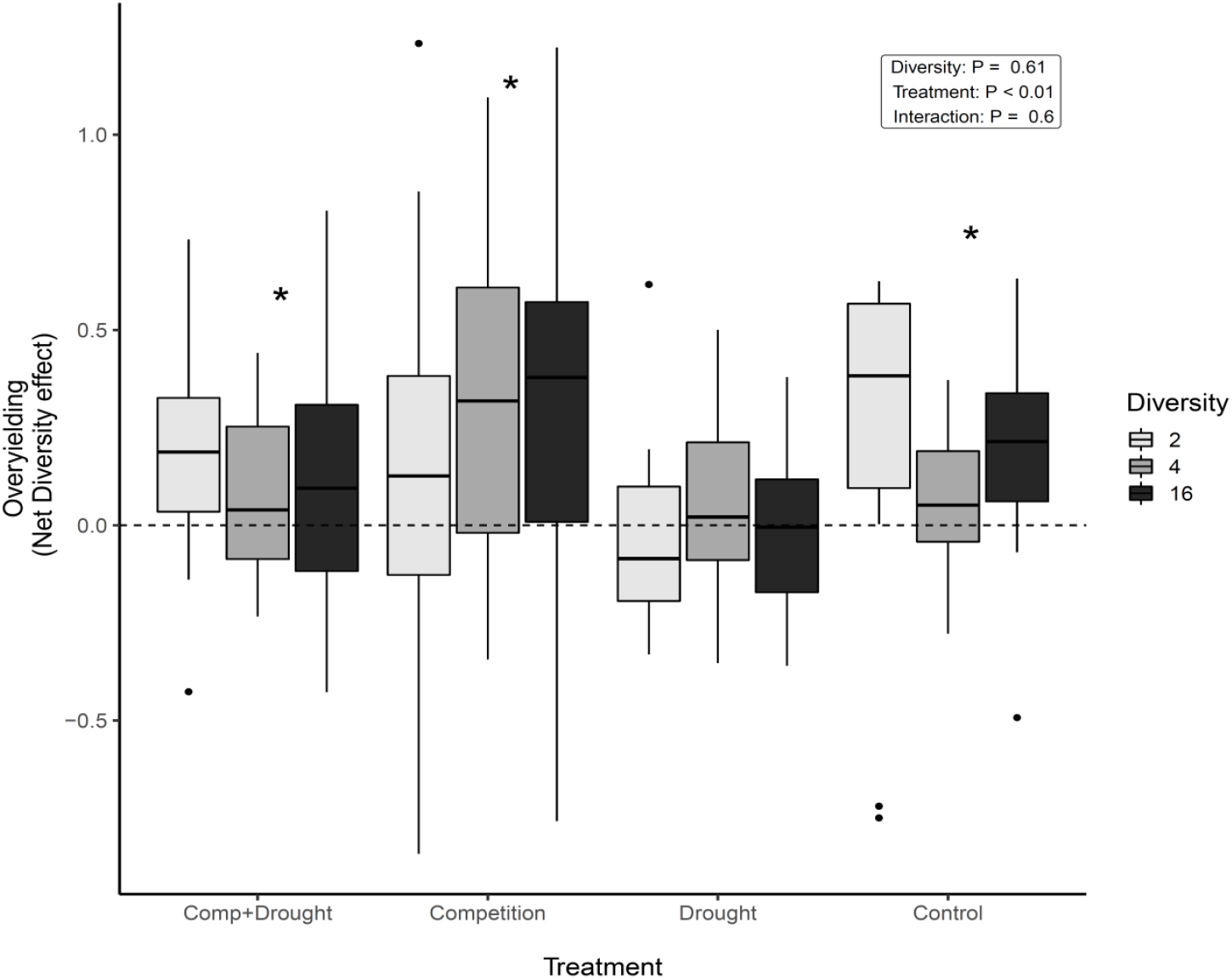
Effect of treatment and number of TE lines on overyielding of the mixtures (how better mixtures perform than monocultures) calculated as net diversity effect following the formula of Loreau and Hector, 2001 (i.e. for a given mixture, it is the difference between the observed productivity and the expected productivity, which is the productivity of the different TE lines that comprise the population in the corresponding monoculture). Positive values imply that the observed productivity of the mixture is higher than expected from monocultures. Asterisks indicate significant positive overyielding of the treatments (significantly different from zero). The p value of the effect of each of the factor are shown in the box.

**Table S1:** Summary of performances of all the models conducted to predict Arabidopsis productivity and competitor’s performance based on treatment, population diversity measured as number of lines (Model 1) or functional diversity (Model 3) and the presence of specific TE lines (Model 2) or population trait means (Model 4) and their respective combinations (Model 5 & 6). The best model is the result of forward selection of predictors based on the lowest AIC; thus, variables are selected if they increase model fit and no matter whether they have significant effect. For each model, significant effects of the selected variables are shown in bold, and the colour indicates the direction of the effect (positive in blue, negative in red). The R^2^ represents the variation explained by the selected variables whereas the *Res*. *df* represents the residuals degrees of freedom.

| Model | Focus | Candidate variable(s) | Arabidopsis productivity |  |  |  | Competitors performance |  |  |  |
| --- | --- | --- | --- | --- | --- | --- | --- | --- | --- | --- |
|  |  |  | Selected variables | AIC | Res. df | R<sup>2</sup> | Selected variables | AIC | Res. df | R<sup>2</sup> |
| 1 | Number of lines | Treatments x log(Diversity) | Treatments + **log(Diversity)** + Treat:Diversity(ns) | 135.1 | 312 | 0.779 | Treatments | 174.8 | 158 | 0.188 |
| 2 | Functional diversity | Treatments x Rao | Treatments+**Rao**+ Treat:Diversity(ns) | 134.6 | 312 | 0.779 | Treatments + **Rao** | 171.2 | 157 | 0.212 |
| 3 | Presence of specific lines | Forward selection from all lines: +L1 +L2 + (...) +L20 | Treatments + **L8(ns)** + **L15** + **L18** | 130.6 | 313 | 0.781 | Treatments + **L1(ns)** + **L2** + **L5** + **L7(ns)** + **L11** + **L15(ns)** | 162.1 | 152 | 0.277 |
| 4 | Population functional trait means | Forward selection from populations' trait means (SLA, LDMC, SRL...) | Treatments + **SLA** | 137.6 | 315 | 0.775 | Treatments | 174.8 | 158 | 0.188 |
| 5 | Number of lines + Presence of specific lines | Candidate variables from Model 1 & 3 | Treatments + **log(Diversity)** + **L5(ns)** + **L8** + **L15** + **L20(ns)** + Treat:Diversity(ns) | 126.9 | 308 | 0.787 | Treatments + **log(Diversity)(ns)** + **L1(ns)** + **L2** + **L5** + **L7(ns)** + **L11** + **L15(ns)** + **L16(ns)** + **L19(ns)** | 159.2 | 149 | 0.302 |
| 6 | Functional diversity + Population functional trait means | Candidate variables from Model 2 & 4 | Treatments + **Rao** + **SLA** | 126.0 | 314 | 0.784 | Treatments + **Rao** + **SLA(ns)** | 171.0 | 156 | 0.217 |

